# Interactive effects of land use change and climate drive the decline of a threatened boreal forest orchid in Finland

**DOI:** 10.64898/2026.10.12.758414

**Authors:** S. H. M. Koivusaari, M. Luoto, J. Aalto, K. Aapala, M. H. Hällfors, R. K. Heikkinen, T. Laurinen, A. L. K. Mattila

## Abstract

Land use and climate change are two of the greatest threats to biodiversity globally. However, their interactive effects remain poorly understood. In the boreal forest biome, where species are experiencing increasing pressures from forest management and climate change, improved understanding of these effects is urgently needed. Here, we assess the impacts of forest loss and both baseline and changing late spring climatic conditions on the extinction probability of a threatened boreal forest orchid, *Calypso bulbosa*. We use a unique dataset consisting of 1,006 *C. bulbosa* occurrence sites located in Finland, historical aerial images, and climatic data spanning the last 60 years. Our results reveal that forest loss, at the local rather than landscape-scale, and both the site-specific baseline climatic conditions and their change over time influence the extinction probability of *C. bulbosa.* These effects were mainly interactive with greater temporal increase in rainfall ameliorating and greater baseline rainfall exacerbating the negative effect of forest loss on the species’ persistence. Furthermore, greater baseline snow depth ameliorated this negative effect. We also found some statistical support for independent effects of baseline thermal sum and its change over time, but their ecological relevance was interpreted as minor. Overall, our results align with the expectation that forest loss can modify the exposure of forest-dependent species to climatic conditions, thus altering their vulnerability to climate change.

## 1. INTRODUCTION

Biodiversity is declining at an alarming rate. This decline has been mainly attributed to the loss, fragmentation, and degradation of habitats caused by land use change (Jaureguiberry et al., 2022; Semenchuk et al., 2022). However, climate change is increasingly contributing to the loss of biodiversity by exposing species to novel climatic conditions at a pace that exceeds their capacity to adjust, adapt, and disperse (Mattila et al., 2024; Román-Palacios & Wiens, 2020; Thomas et al., 2004). While land use and climate change can contribute to biodiversity decline independent of one another (individual and additive effects), they may also have interactive effects (Oliver & Morecroft, 2014). For instance, land use change can exacerbate the negative impacts of climate change on species persistence by reducing their ability to track suitable climatic conditions through dispersal. Although the interactive effects of land use and climate change are increasingly assessed (Auffret & Svenning, 2022; Northrup et al., 2019; Suppula et al., 2023), the prevalence of such effects, as well as the mechanisms by which they operate in different ecosystems, remain poorly understood.

The boreal forest biome is experiencing increasing pressures from both land use and climate change. The area and quality of natural boreal forest is rapidly declining, mainly due to the expansion and intensification of commercial logging (Burton et al., 2010; Gauthier et al., 2015). Additionally, the boreal forest biome is warming two to four times faster than the global average due to its high-latitude location (Rantanen et al., 2022). Importantly, not only are mean temperatures rising, but climate change is also altering precipitation and snow patterns (Aalto et al., 2023). Furthermore, extreme weather events are increasing in frequency and intensity (Aalto et al., 2026; Rao et al. 2023). These land use and climatic changes can interact with one another in a distinct way. Forest management activities, such as thinning and clear-cutting, are known to diminish the microclimatic buffering capacity of boreal forests (Chen et al., 1993; Starck et al., 2025, 2026). Reductions in the forests’ microclimatic buffering capacity, in turn, can expose species inhabiting them to more severe impacts of climate change compared to those that they would otherwise experience (De Frenne et al., 2019; Geiger, 1965; Zellweger et al., 2020).

Here, our aim is to assess the individual, additive, and interactive effects of land use and climate change on the decline of a boreal forest orchid, *Calypso bulbosa* (L.) Oakes. So far, the decline of *C. bulbosa* has been mainly linked to habitat loss and degradation caused by forest management and construction (Hyvärinen et al. 2019). However, climate change may increasingly impact the species’ survival, especially if forestry practices diminish microclimatic buffering. For instance, increasing temperatures, declining snow cover, and night frosts in spring may negatively influence *C. bulbosa* by exposing the species to drought and frost damage (Chen et al., 1993; Hu et al., 2023; Örlander, 1993; Simons et al., 2010; Wu et al., 2025; Yan et al., 2025). Increases in precipitation, on the other hand, may be beneficial for *C. bulbosa* as they may improve the moisture conditions (Koivusaari et al., 2026). However, extensive increases in rainfall could lead to root suffocation and thus influence the species negatively (Hirabayashi et al., 2013; Irfan et al., 2010; Zhang et al., 2025).

The degree at which land use and climate change influence *C. bulbosa* may depend on the spatial scale at which land use change impacts manifest, as well as on the baseline climatic conditions in the area. Species with low dispersal ability, such as *C. bulbosa*, may be more vulnerable to changes that occur in their close vicinity than those with higher dispersal ability and thus greater potential to rescue populations and recolonize after local extinctions (Brown & Kodric-Brown, 1977; Saccheri et al., 1998), although the evidence for such differences between weak and strong dispersers remains limited (Martin et al., 2023). Furthermore, climatic changes may have more severe impacts on species at sites where baseline climatic conditions are already close to their environmental tolerance limits (Deutsch et al., 2008). Conversely, at sites where climatic conditions are close to the species’ environmental optimum, populations may remain stable despite equally pronounced climatic changes (Deutsch et al., 2008).

To assess the interactive impacts of land use change and climate (baseline and change) on the decline of *C. bulbosa*, we use a unique dataset consisting of 1006 *C. bulbosa* occurrence sites located in Finland, historical aerial images, and climatic data spanning the last 60 years. We expect forest loss to increase the extinction probability of *C. bulbosa* while higher baseline and increasing thermal accumulation and lower baseline and decreasing snow during early spring would increase, and higher baseline and increasing rainfall decrease extinction probability. In addition to these individual effects, we anticipate forest loss to impact the microclimatic buffering capacity of the forest, manifesting as interactive effects of land use and climate on the extinction probability of *C. bulbosa*. In particular, we expect greater baseline and increase in thermal accumulation and lower baseline and greater decrease snow cover to have exacerbating interactive effects on extinction probability together with land use, while greater baseline and increased rainfall would have an ameliorating interactive effect, provided that the conditions are not becoming overly wet. Finally, we expect forest loss at the site-scale to have a more important impact on *C. bulbosa* in comparison to forest loss at the landscape-scale.

## 2. METHODS

### 2.1 Study species

*Calypso bulbosa* (L.) Oakes (Orchidaceae) (Fig. 1A) is a perennial herb with a circumboreal distribution (Hultén & Fries 1986). In Finland, it usually grows in shady old-growth coniferous forests with high soil calcium concentration (Abeli et al., 2013; Tuomi et al., 2015). The species flowers soon after snow melt, usually in late May or the beginning of June (Alexandersson & Ågren, 2000). The reproductive stage of the species forms one flower which can produce thousands of small, wind-dispersed seeds (Alexandersson & Ågren, 2000; Boyden, 1982). Like other orchids, *C. bulbosa* relies on relatively complex biotic interactions: it obtains organic carbon from its mycorrhizal partner (Currah et al., 1988; Hadley, 1982) and is pollinated by bumblebee queens (Ackerman, 1981; Alexandersson & Ågren, 1996; Boyden, 1982; Mosquin, 1970).

**Figure 1.**
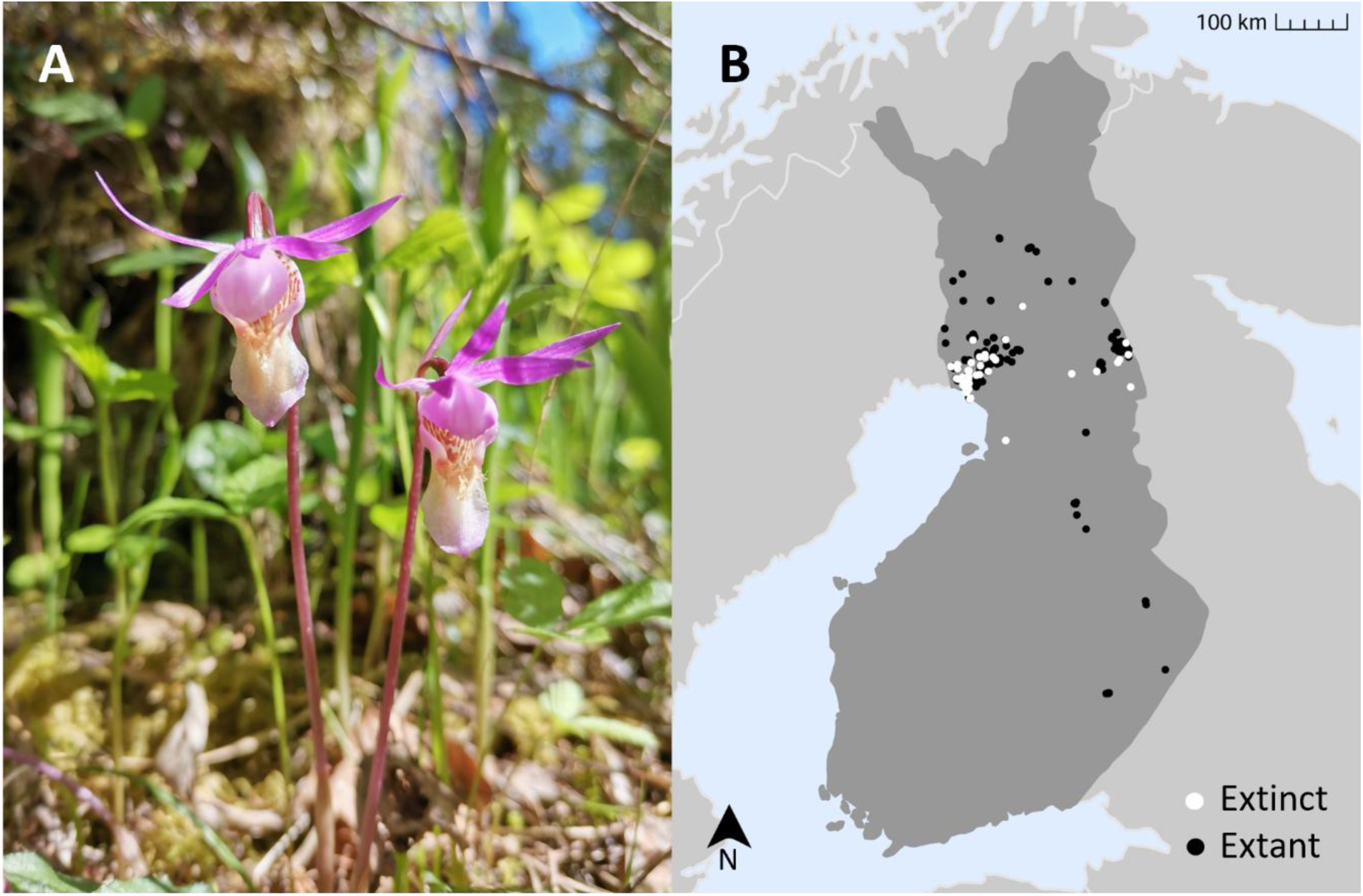
Illustration of the study species and sites. Panel A shows a picture of the study species *C. bulbosa* (photograph by S. Koivusaari). Panel B shows the locations of the extant (black) and extinct (white) study sites in Finland.

### 2.2 Data

#### 2.2.1 Species occurrence data

We obtained species occurrence data from the databases of the Finnish Environment Institute (HERTTA) and Parks & Wildlife Finland (LajiGIS, Silvia). The data are available from the Finnish Biodiversity Information Facility (FinBIF, 2023; collection ID MX.40083). The species data were downloaded in 2020 and 2023. The data consisted of occurrence records based on visitations at known occurrence sites of *C. bulbosa* across Finland. At each visit, the status of the occurrence site was recorded to be either extant, extinct, or uncertain. An uncertain status was assigned when an occurrence site that was previously confirmed as extant was not detected during the current field visit, and the available evidence was insufficient to conclude that the species had become locally extinct. If an occurrence site received a different status in different years, we assigned it according to the most recently recorded status. We omitted occurrence sites for which the status remained uncertain.

For the extinct sites, we also determined the year when the extinction likely occurred. If an occurrence site had been recorded as extinct in the species data, the year of extinction was usually reported at an accuracy of one to two decades. For 19 sites, however, no year of extinction was provided. For these sites, we determined a window of extinction based on the notes made by the observers. These notes often contained information on when the species was last observed at the occurrence site (even though these intermediate observations had not always been added as separate visits to the database), and when it was first observed as extinct. After this, we assigned the year of extinction by either assigning a mid-decade year for decadal-level data (e.g. if the occurrence site was recorded to have gone extinct in the 1990’s, we set the extinction year to 1995) or a mean year for extinction time window information (e.g. if the extinction window was 2004-2010, we set the extinction year to 2007). As our climatic data started in 1961 (see Section 2.2.4) and to allow estimating climatic trends over a minimum of 15 years, we excluded occurrence sites for which the last observation year (if extant) and extinction year (if extinct) occurred prior to 1975.

To improve the spatial resolution of the data, we excluded records with coordinate accuracy of >10 meters. Furthermore, to avoid pseudoreplication, we retained only one occurrence site per 20 x 20 m cell. This spatial filtering was done simultaneously for extant and extinct sites, because all the 20 x 20 m cells contained either occurrences deemed as extant or extinct, and not both types of states. After this data processing, the data contained 2575 unique occurrence sites, of which 84 were recorded as extinct and 2491 extant.

#### 2.2.2 Connectivity data

To capture the spatial structure of the studied *C. bulbosa* occurrence sites, we used the measure of population connectivity described by Hanski (1994). This measure accounts for distance from each occurrence site to all potential source populations and the population size of all potential source sites, and can be defined through the following equation:

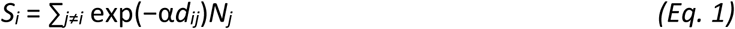

where *S_i_* is a measure of connectivity for population ‘*i*’ to other potential source populations ‘*j*’; α scales the effect of distance to migration; *d_ij_* is the distance (in kilometers) from patch *i* to *j*; and *N_j_* is the population size of the potential source population.

The method assumes that the data contains information on all populations of the species. However, to calculate the measure of connectivity, we used occurrence data containing 2575 unique occurrence sites filtered based on the rules described in Section 2.2.1. In these data, abundance was recorded at least once for 1901 of the sites, allowing us to calculate mean abundance across years per occurrence site to indicate population size. Both the unit and format of abundance data varied across records. For most records, the unit was the count of individuals or sprouts, but for some records, it was defined as the total individual count or the count of flowering individuals or sprouts/tufts. For most records the abundance value was given as a numerical value, but for 16 sites, abundance was given as a verbal description, such as “multiple” or “a few”. Furthermore, in 34 sites, abundance was recorded as 0 at all visits. In cases where the unit was something else than individuals or sprouts and the abundance value was reported verbally, we assigned a value for abundance based on the description (e.g. a few was translated to 5 individuals). To handle occurrence sites with missing or zero mean abundance, we used two alternative methods: we assigned these occurrence sites 1) a random mean abundance value within the range of the most common mean abundance values (within 2.5 and 97.5 % quantiles), and 2) a mean abundance value of 1. Using mean abundance data constructed with these two alternative approaches, we calculated connectivity according to Equation 1, with α set to 1 based on the estimation of dispersal ability of *C. bulbosa* (Lososová et al., 2023) and compared the results obtained via both approaches (Section 3.2).

#### 2.2.3 Land use change data

To assess land use changes in the study sites, we used historical aerial images provided by the National Land Survey of Finland. For each year for which an aerial image was available for an occurrence site, we visually estimated the percentage of area covered by older forest (i.e. forest with continuous canopy and no signs of management, e.g. forestry machinery tracks) out of the total land area within a 250 m radius from the occurrence site. We then calculated the change in the older forest cover for each time interval (i.e. % mature forest in year_i_ -% mature forest in year_i-1_). Finally, we summed the differences across all time intervals to obtain a measure of cumulative change in the forest cover. For the extant sites, we included information extracted from aerial images taken during the years before the occurrence site had been visited for the last time (varying between 1987-2022) and from the next year after the last visit for which aerial images were available. For the extinct sites, we included information from the years before the occurrence site was deemed extinct (ranging between 1975-2019; see Section 2.2.1 on how the extinction year was determined), and from the next year after the extinction event for which aerial images were available. Based on this filtering process, two to 14 aerial images per occurrence site were included in the analyses.

In addition to the percentage of area covered by older forest within the 250 m buffer area, we recorded information on whether the exact location of the occurrence site was covered by continuous forest. This information was also collected based on visual inspection of aerial images. We then estimated linear change in the presence of mature forest by calculating Sen’s slope (Sen, 1968) separately for each occurrence site using the *sens.slope* function in the ‘trend’ R package (Pohlert, 2023). To make sure that the slope captures the potential changes that have led to the extinction of the occurrence sites as accurately as possible, we included only information from 1) the previous available year before the final observation year (if extant) or the year of extinction the recorded event (if extinct), and 2) the next year after the final observation year or year of extinction event for which aerial images were available. We classified the slope values into four classes based on how suitable the class was expected to be for *C. bulbosa* (in the order from least suitable): 1) loss of forest; 2) no forest over the examined temporal span; 3) gain of forest; and 4) continuous forest over the examined temporal span. We assumed class 1 to be the least suitable for this species, given that it reflects a disturbance. Class 2 was expected to be more suitable given that, in some cases, it is possible that the populations occur in less suitable, undisturbed conditions. Class 3 was assumed to be more suitable for the species, given that mature forest would have developed, while class 4 was assumed to describe the most suitable conditions as it reflected situations where forest had been continuously present at the site.

We assessed aerial images for all extinct occurrence sites for which there was aerial image data available, and for a sample of extant sites. The sample was based on a random selection of locations, with priority given to locations with <80-year-old forest cover to reduce age-related sampling bias and improve the geographic representation of sites across the two main study areas. As a result, we obtained land use change data for 83 extinct and 924 extant sites, totaling 1006 sites.

**Figure 2.**
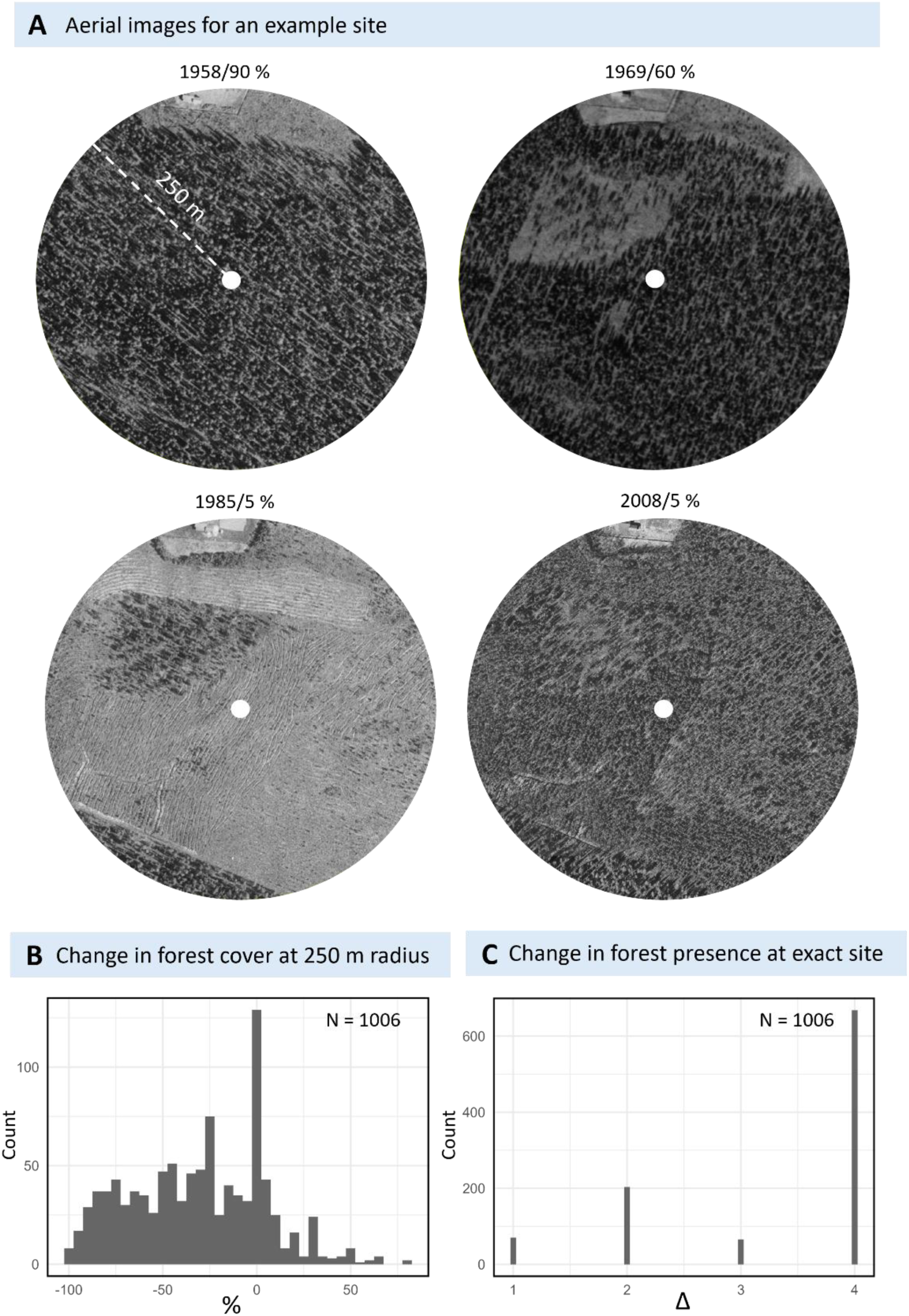
Illustration of the land use change data. Panel A shows an example of an aerial image series used to assess changes in land use. The study occurrence site is shown with a white point in the center of each aerial image. The estimated % of forest cover is given after the year for each image. Panel B shows values obtained for cumulative change in forest cover within a 250 m buffer area (%). Panel C shows linear change in the presence of forest in the exact location of the study occurrence site (Δ). Class 1 refers to the loss of forest, class 2 to no forest cover at all, class 3 to the gain of forest, and class 4 to continuous forest cover.

#### 2.2.4 Climate data

To assess changes in the climatic conditions, we used daily gridded climate data at 1 x 1 km spatial resolution provided by the Finnish Meteorological Institute (Aalto et al., 2016). We selected three variables to use in our analyses: 1) rainfall (mm); 2) growing degree days above 5 °C (GDD5; sum of temperatures exceeding +5 °C); and 3) mean snow depth (cm). Rainfall and GDD5 were calculated annually for 1961–2019 as the mean of accumulated rainfall and degree days over April-June, representing average late spring-month rainfall and GDD5, respectively. Mean snow depth, in turn, was calculated annually for 1961-2020 as the mean snow depth during April and May. Here, June was excluded to avoid bias caused by the large number of zero-values (the average site had no snow cover in June during 59 of the 60 years). Overall, for the climatic variables, these late-spring months were considered the most relevant for the ecology of the study species, given that it flowers during spring and early summer. To match the spatial resolution of climatic data to that of the species data, we resampled the climatic data to a 20 x 20 m resolution using a bilinear interpolation method and the function *project* in ‘terra’ R package (Hijmans, 2025).

We estimated linear trends in the climate variables over 1961-2019 (2020 for snow depth) by calculating Sen’s slope (Sen, 1968) separately for each of the 1007 occurrence sites for which we also had land use change data, using the *sens.slope* function in the ‘trend’ R package (Pohlert, 2023). For the extant sites, Sen’s slope was calculated starting from the first year of available climatic data (1961) to the final observation year, whereas for the extinct sites, it was calculated until the year when the population was deemed extinct.

Similarly for forest cover data, the last observation year varied between 1987-2022 for extant occurrence sites while the extinction designation varied between 1975-2019. The mean number of years used to estimate climatic trends across all occurrence sites was 50.84 years, while the mean across extant and extinct occurrence sites was 40.01 and 51.21 years, respectively. We removed one occurrence site where the change in the amount of rainfall fell clearly outside the range of all other values (i.e. was an outlier or an error in the data), totaling 82 extinct and 924 extant occurrence sites.

In addition to estimating change in the climatic variables over time, we calculated the occurrence site-specific average value of each climatic variable across the first 30 years of the study period (1961-1990). This was done to allow controlling for occurrence site-specific baseline climatic conditions in subsequent analyses and thus to make sure the potential effect observed from the slope of the linear change would in fact reflect the impact of change in climatic conditions rather than differences in baseline climatic conditions between the sites.

**Figure 3.**
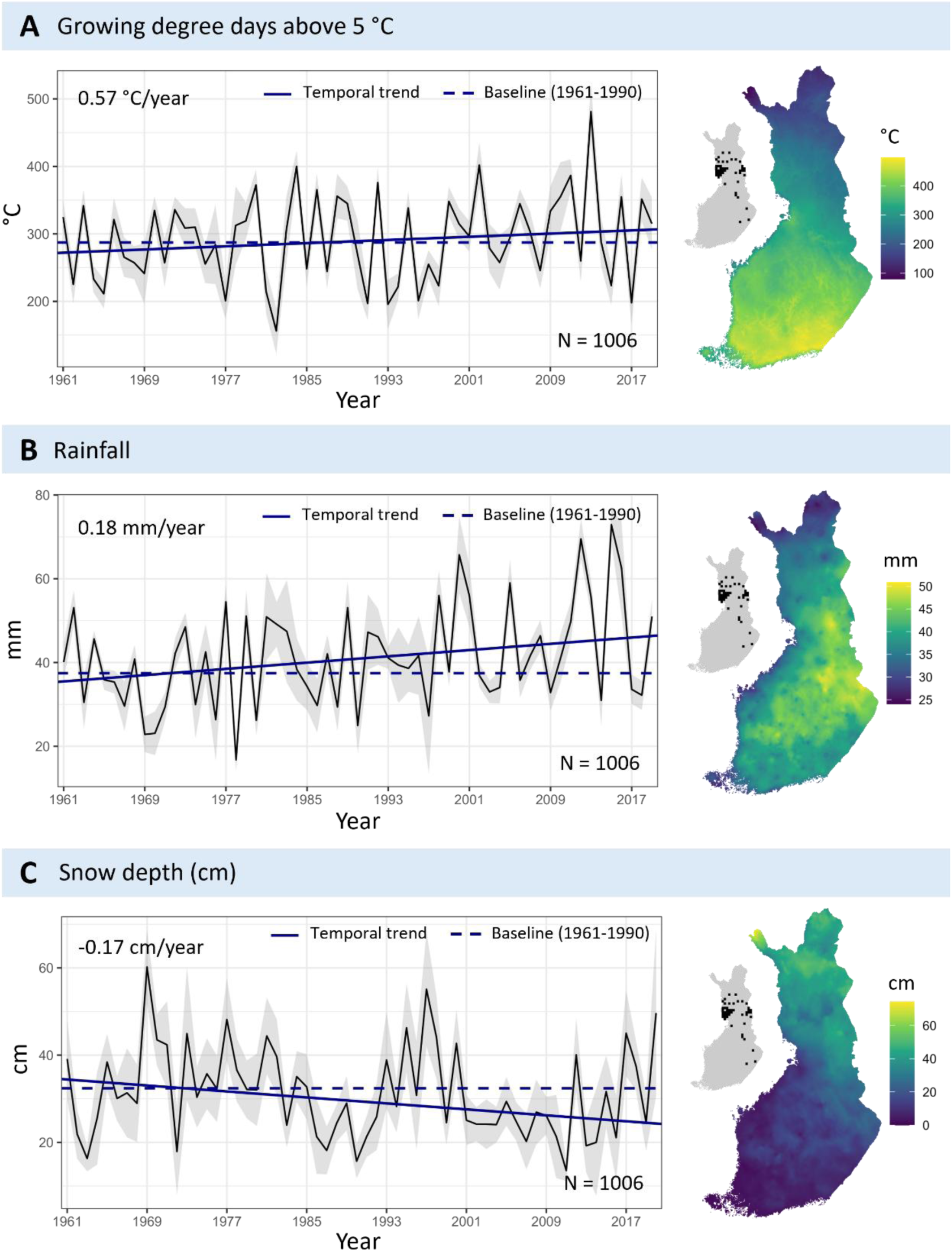
Illustration of the climatic data. In each panel, the black line shows temporal fluctuations in climatic conditions across the study period, averaged across all study sites, while the grey areas show standard deviation in climatic conditions across all study sites. The blue dashed lines show baseline climatic conditions across the first 30 years of the study period and the blue solid line shows the linear temporal trend, both averaged across all study sites. The temporal span used to calculate linear change was adjusted per study occurrence site (see Section 2.2.4). The values in the panels show change in each climatic variable per year across study sites. The maps in the right side of each panel show the spatial distribution of climatic conditions in Finland, averaged across the first 30 years in the climatic data (1961-1990). The locations of the study occurrence sites are shown as 20 x 20 km grid squares in the smaller maps.

### 2.3 Statistical analyses

All analyses were conducted in R version 4.5.1 (R Core team 2025).

#### 2.3.1 Model fitting

We used Generalized Linear Models (GLMs) with a binomial error distribution and logit link function to model the binary status (extinct = 1, extant = 0) of the 1006 occurrence sites of *C. bulbosa*. The models were implemented with the *glm* function from the ‘stats’ R package (R Core Team, 2025). To assess the additive and interactive effects of land use and climate on the extinction probability of *C. bulbosa*, we fitted four different models that varied in their main predictors: 1) land use change only; 2) climate only; 3) land use and climate; and 4) land use change, climate, and their interactions (Table 1). The climate component always included both baseline climatic conditions and changes in them. In all models, we included connectivity to capture the spatial structure of the data, and in model 4. Two alternative sets of models were fitted using connectivity data calculated with two methods (see Section 2.2.2). The fit of the models with different main predictors, as well as different connectivity data were compared using AIC and Nagelkerke R^2^ values. Given the large range of connectivity values in comparison to the other explanatory variables, connectivity was square-root transformed prior to modeling. In model 4 (the interaction model), we initially included all possible interactions between land use change and climate, and land use change and connectivity, and climate and connectivity. Then, we used a backward-selection approach to exclude non-significant (p > 0.05) interactions.

**Table 1.** The structure of the models and their usage in terms of interpreting the impacts of land use and climate on the extinction probability of *C. bulbosa*. In all models, connectivity, estimated with two alternative approaches, was included to capture the spatial structure of the data. LUC = land use change, C = climate (baseline and change).

| Model Number | Predictors | Usage |
| --- | --- | --- |
| 1 |  | Used to interpret models 3 and 4. |
| 2 | C | Used to interpret models 3 and 4. |
| 3 | LUC + C | Used to assess whether land use change and climate (baseline and change) have additive effects, i.e. whether including both land use change and climate improves model fit compared to models including only one of them. This also implied assessing whether their effects remained significant when controlling for the other.<br>This model was also used to interpret model 4. |
| 4 | LUC + C + LUC:C | Used to assess whether land use change and climate (baseline and change) have interactive effects, i.e. whether the impacts (or explanatory power of the model) of land use change or climate is stronger or weaker under different land use change and climate conditions. |

#### 2.3.2 Model assessment

We assessed multicollinearity of our predictors using the Variance Inflation Factor (VIF). The VIF values, which were calculated based on model 3 (see Table 1) to exclude influence of the interaction terms, were generally low (below 1.7), but for baseline GDD5 and baseline rainfall, the VIF values indicated moderate multicollinearity (VIF = 4.02 and 5.48, respectively using connectivity approach 1 and 5.07 and 6.13, respectively using connectivity approach 2). This multicollinearity was taken into account with cautious interpretation of the model estimates for these two variables. To assess residual spatial autocorrelation, we used the *testSpatialAutocorrelation* function from the ‘DHARMa’ R package (Hartig, 2024) with residuals simulated with the *simulateResiduals* function. The test, performed on model 4, did not reveal spatial autocorrelation in the residuals (Moran’s I = 0.01, p = 0.43 using connectivity approach 1 and Moran’s I = 0.01, p = 0.52 using connectivity approach 2). To assess the performance of the models, we used the Receiver Operating Characteristic (ROC) curve and Area Under the Curve (AUC) value produced with the *roc* and *auc* functions from the ‘pROC’ R package (Robin et al. 2011), as well as QQ-plots and residuals vs. fitted values plots (Fig. S1) created with the *plot* function and simulated residuals from the *‘*DHARMa’ R package. The ROC curve and AUC values indicated excellent discriminatory performance (Fig. S2). The QQ-plots and residuals vs. fitted values plots did not indicate deviations from the normality of residuals and homogeneity of variances assumptions (Fig. S2). We used the *ggpredict* function from the ‘ggeffects’ R package (Lüdecke, 2018) to plot partial effects of all variables used in the best performing models (i.e. those with the lowest AIC and highest R^2^ value).

## 3. RESULTS

### 3.1 Land use and climate change

The cover of forest in the landscape surrounding the occurrence sites decreased (within the timespan that was adjusted per site), on average, by 32 % (Table 2). Across sites, changes ranged from a 100% decrease to an 80% increase. The categorical changes in the presence of forest cover at the exact location of the occurrence site varied between loss of forest cover, no change (forest cover was either continuously present or absent), and gain of forest cover. Specifically, in 6.95 % of sites, forest had disappeared, in 20.18 % of sites forest cover was not present during the entire study period, in 6.46 % of sites forest had recovered during the study period, and in 66.40 % of sites it was continuously present. Rainfall increased on average by 0.18 mm per year (−0.74 - +0.43 mm per year), while snow depth decreased on average by 0.17 cm per year (−0.43 - +0.31 cm per year), and GDD5 increased on average by 0.57 days per year (−0.99 - +1.77 °C per year).

**Table 2.** Summary statistics of the explanatory variables. Status = the status of the occurrence site (extant/extinct), CMP = change in mature forest presence, CMC = change in mature forest cover, CRF = change in rainfall, CGDD = change in GDD5, CSD = change in snow depth, MRF = baseline rainfall, MGDD = baseline GDD5, MSD = baseline snow depth, CON1 = connectivity method 1, CON2 = connectivity method 2.

| Variable | Min | Mean | Median | Max | SD |
| --- | --- | --- | --- | --- | --- |
| Status | 0 | 0.08 | 0 | 1 | 0.27 |
| CMP | 1 | 3.32 | 4 | 4 | 1.02 |
| CMC | -100 | -31.53 | -30 | 80 | 35.65 |
| CRF | -0.14 | 0.18 | 0.20 | 0.43 | 0.08 |
| CSD | -0.43 | -0.17 | -0.17 | 0.31 | 0.08 |
| CGDD | -0.99 | 0.57 | 0.60 | 1.77 | 0.29 |
| BRF | 29.56 | 37.36 | 36.02 | 47.94 | 3.32 |
| BGDD | 219.34 | 291.12 | 293.71 | 435.79 | 31.81 |
| BSD | 10.16 | 31.62 | 33.92 | 45.99 | 6.56 |
| CON1 | 0 | 1808.68 | 969.77 | 6290.77 | 1909.68 |
| CON2 | 0 | 432.08 | 359.41 | 3273.91 | 437.8 |

### 3.2 Comparison of models fitted with different connectivity-calculation approaches

Based on a comparison of AIC values, models fitted using the two different approaches to estimate connectivity values did not significantly differ in their performance (ΔAIC < 2; Table 3; Table S1). Similarly, there were only minor differences in the amount of variation explained by the models (R^2^ = 0.58 and R^2^ = 0.57 for approach 1 and 2, respectively). However, as we only wanted to include one connectivity estimate, we chose to use approach 2 given its slightly better performance in terms of AIC (Table 3) and our assumption that it was more ecologically meaningful than approach 1, which was more likely to produce unrealistically high connectivity values given that a random mean abundance value was given to all sites with 0 or NA mean abundance.

**Table 3.** Model comparison results for two sets of four models fitted using connectivity data either estimated with approach 1 or 2. In connectivity estimation approach 1, random mean abundance values were given to occurrence sites with NA and 0 mean abundances, and in connectivity estimation approach2, a mean abundance value of 1 was given to these sites. LUC = land use change, C = climate (baseline and change).

| Connectivity method | Model | R<sup>2</sup> | AIC | ΔAIC |
| --- | --- | --- | --- | --- |
| 1 | LUC | 0.27 | 449.38 | 135.27 |
| 1 | C | 0.25 | 467.70 | 153.59 |
| 1 | LUC + C | 0.48 | 355.69 | 41.58 |
| 1 | LUC + C + LUC:C | 0.58 | 314.11 | 0.00 |
| 2 | LUC | 0.30 | 434.55 | 122.19 |
| 2 | C | 0.33 | 428.43 | 116.07 |
| 2 | LUC + C | 0.50 | 345.95 | 33.59 |
| 2 | LUC + C + LUC:C | 0.57 | 312.36 | 0.00 |

**Table 4.** Summary statistics for the four models used to understand the independent, additive, and interactive effects of land use and climate on the extinction probability of *C. bulbosa*. LUC = land use change, C = climate (baseline and change). CMP = change in mature forest presence, CMC = change in mature forest cover, CRF = change in rainfall, MRF = baseline rainfall, CGDD = change in GDD5, MGDD = baseline GDD5, CSD = change in snow depth, MSD = baseline snow depth, CON = connectivity.

| Model | Term | Estimate | Std. Error | Statistic | p-value |
| --- | --- | --- | --- | --- | --- |
| LUC | (Intercept) | 1.48 | 0.43 | 3.46 | <0.01 |
|  | CMP | -1.06 | 0.13 | -7.92 | <0.001 |
|  | CMC | 0.00 | 0.00 | -0.06 | >0.1 |
|  | CON | -0.07 | 0.02 | -3.82 | <0.001 |
| C | (Intercept) | 8.00 | 3.13 | 2.55 | 0.011 |
|  | CRF | -10.64 | 1.74 | -6.10 | <0.001 |
|  | CGDD | -1.10 | 0.45 | -2.47 | <0.05 |
|  | CSD | 2.28 | 1.79 | 1.28 | >0.1 |
|  | BRF | -0.23 | 0.06 | -3.85 | <0.001 |
|  | BGDD | 0.01 | 0.01 | 0.87 | >0.1 |
|  | BSD | 0.01 | 0.05 | 0.16 | >0.1 |
|  | CON | -0.11 | 0.02 | -6.19 | <0.001 |
| LUC + C | (Intercept) | 8.06 | 3.81 | 2.12 | <0.05 |
|  | CMP | -1.28 | 0.17 | -7.56 | <0.001 |
|  | CMC | 0.00 | 0.00 | -0.02 | >0.1 |
|  | CRF | -14.11 | 2.17 | -6.50 | <0.001 |
|  | CGDD | -0.54 | 0.48 | -1.11 | >0.1 |
|  | CSD | 2.79 | 1.83 | 1.52 | >0.1 |
|  | BRF | -0.10 | 0.07 | -1.42 | >0.1 |
|  | BGDD | 0.01 | 0.01 | 0.65 | >0.1 |
|  | BSD | -0.04 | 0.06 | -0.70 | >0.1 |
|  | CON | -0.07 | 0.02 | -3.00 | <0.01 |
| LUC + C + LUC:C | (Intercept) | -18.17 | 9.44 | -1.92 | <0.1 |
|  | CMP | 7.11 | 2.44 | 2.92 | <0.01 |
|  | CMC | 0.00 | 0.00 | -0.26 | >0.1 |
|  | CRF | 3.22 | 4.42 | 0.73 | >0.1 |
|  | CGDD | -1.24 | 0.53 | -2.35 | <0.05 |
|  | CSD | 1.12 | 1.80 | 0.62 | >0.1 |
|  | BRF | 0.75 | 0.27 | 2.79 | <0.01 |
|  | BGDD | 0.03 | 0.01 | 2.37 | <0.05 |
|  | BSD | -0.51 | 0.14 | -3.71 | <0.001 |
|  | CON | -0.50 | 0.13 | -3.98 | <0.001 |
|  | CMP:CRF | -8.60 | 2.09 | -4.11 | <0.001 |
|  | CMP:BRF | -0.30 | 0.09 | -3.45 | <0.01 |
|  | CMP:BSD | 0.13 | 0.04 | 2.97 | <0.01 |
|  | BSD:CON | 0.01 | 0.00 | 3.51 | <0.001 |

### 3.3 Comparison of models with different main predictor set

The model that included both land use change and climate (baseline and change), in addition to connectivity, explained a greater proportion of variation in extinction probability (R^2^ = 0.50; Table 3) than the models including only land use change (R^2^ = 0.30) or climate (R^2^ = 33), in addition to connectivity (Table 2). However, extinction probability was best explained by the model that included also the significant interaction between land use change, climate, and connectivity (R^2^ = 0.57). This model performed the best also based on the comparison of AIC values (ΔAIC < 2; Table 3). Thus, this model was used for inference.

### 3.4 The effects of land use change, climate, and connectivity

The baseline growing degree days above 5 °C (GDD5), as well as temporal change in GDD5 were associated with extinction probability on their own. Extinction probability was greatest at sites characterized by higher baseline GDD5 and greater decrease in GDD5 (Est. = 0.03, p < 0.05; and Est. = −1.24, p < 0.05 for baseline and change in GDD5, respectively; Table 3). Yet, the effect of baseline GDD5 should be inferred with caution due to the moderate multicollinearity it showed in the models. Interactive effects, in turn, were found among forest loss, changes in rainfall, baseline rainfall, and baseline snow depth. Extinction probability was greatest at occurrence sites that had experienced forest loss and the smallest increase in the amount of rainfall (Est. = −8.60, p < 0.001). Furthermore, extinction probability was greatest for occurrence sites that had experienced forest loss and were characterized by the highest baseline rainfall (Est. = −0.30, p < 0.01) and lowest snow depth conditions (Est. = 0.13, p < 0.01). Similarly to baseline GDD5, the effect of baseline rainfall should be inferred with caution due to the moderate multicollinearity it showed in the models. Connectivity was weakly associated with extinction probability in interaction with climate: extinction probability was greatest for occurrence sites that had the highest connectivity and the highest baseline snow depth (Est. = 0.01, p < 0.001). However, in all other models (land use change only, climate only, land use change and climate, Table 1), connectivity decreased extinction probability (Esti. = −0.07, p < 0.001; −0.11, p < 0.001; −0.07, p < 0.01 for LUC, C, and LUC + C models respectively).

**Figure 4.**
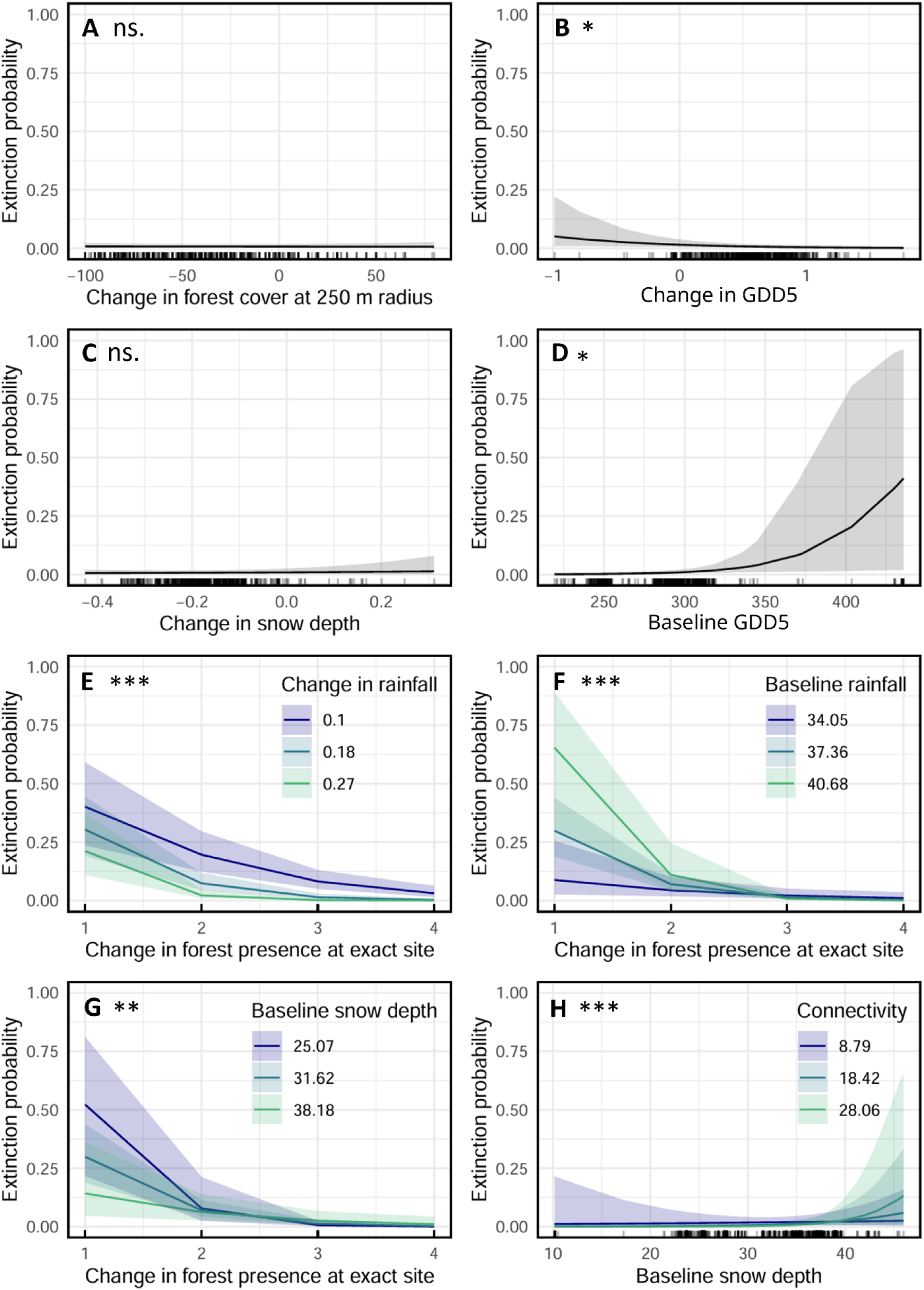
Predicted extinction probabilities across the observed ranges of the land use change, climate change, and connectivity variables. Predictions are based on the model with the lowest AIC and highest R² values, which included all land use change and climate variables (baseline and change), connectivity, and their significant interactions (LUC + C + LUC:C). p < 0.001 = ***, p < 0.01 **, p < 0.5 *. The actual values of the variable in the x-axis are shown as lines in the bottom of the x-axis.

## 4. DISCUSSION

Understanding how land use and climate change jointly influence the persistence of boreal forest species is increasingly important as these pressures intensify simultaneously (Gauthier et al., 2015). Here, we examined their individual, additive, and interactive effects on a declining boreal forest orchid, *Calypso bulbosa*, using a 60-year dataset spanning 1006 occurrence sites across Finland. Our results provide evidence that both baseline climatic conditions and climatic changes modify the effect of forest loss on the persistence of the species. More specifically, we find that the strength of the negative effect of forest loss on the persistence of the species is different at sites with different rainfall and snow conditions. These results highlight that the consequences of habitat loss and degradation cannot, in this case, be understood independently of climatic conditions, and vice versa. These results are consistent with the idea that the loss and deterioration of forests is likely to modify the exposure of species to changing climatic conditions (Zellweger et al., 2020). We also find that the effects of forest loss originate mainly from the local rather than the landscape scale. This result aligns with the general understanding that the effects of habitat loss and degradation are often scale-dependent (Banks-Leite et al., 2020).

### 4.1 Rainfall can ameliorate or exacerbate the effect of forest loss on species persistence

We found that greater temporal increases in rainfall ameliorated the negative effect of forest loss on *C. bulbosa* persistence. This finding was in line with our hypothesis, and could indicate that increasing rainfall compensates for the loss of microclimatic buffering associated with forest loss. Several studies, including one focusing on the occurrence sites of *C. bulbosa* specifically (Koivusaari et al., 2026), have shown that forests can provide different microclimates compared to ambient conditions due to forest structural elements that regulate, e.g., the amount of solar radiation and rainfall penetrating the canopy (Chen et al., 1993; Frey et al., 2016; Meeussen et al., 2021). When the area or quality of a forest is reduced, this buffering capacity can decline and expose forest-inhabiting species to, e.g., drier soil moisture conditions (Chen et al., 1993; Kemppinen et al., 2024; Menge et al., 2023; Senior et al., 2017). Increase in the amount of rainfall, however, could compensate for reductions in the soil moisture level, as suggested by our results. These results highlight that climatic changes may not simply add to the effects of habitat loss but can also modify the ecological consequences of forest degradation. To our knowledge, such interaction has received little attention in the context of plant species persistence in forest habitats.

On the other hand, we observed that greater baseline rainfall level exacerbated the negative effect of forest loss on *C. bulbosa* persistence, although we acknowledge that the effect of baseline rainfall should be inferred with caution given its moderate multicollinearity with the other explanatory variables. These results suggest that although increases in rainfall may, to a certain extent, have a positive effect on *C. bulbosa* populations that experience drought due to forest degradation, there may be a threshold after which exceedingly wet conditions start to reduce the survival of the species by causing flooding and root suffocation (Hirabayashi et al., 2013; Irfan et al., 2010; Zhang et al., 2025). These results highlight that the ecological consequences of the interactive effects of land use and climate change may also depend on how much climate is changing. When assessing the interactive impacts of land use and rainfall, it is also noteworthy that the temporal trend in rainfall varied across the sites of *C. bulbosa*. While most of the sites had experienced increases in rainfall (987), rainfall decreased over time in 19 sites. It is therefore possible that the interactive effects of rainfall and forest loss also depend on the direction of rainfall change. However, our data was not sufficient to assess such bidirectional effects.

Nevertheless, these findings illustrate the complexity of the simultaneous efficacy of land use and climatic changes, and the consequent need for ample data for predicting species responses under such conditions.

### 4.2 Future reductions in snow cover may weaken buffering against the negative impacts of forest loss

We found that deeper baseline snow cover ameliorated the negative impact of forest loss on the persistence of *C. bulbosa*. Snow cover offers insulation for overwintering *C. bulbosa* plants, and the result may indicate a buffering effect of deeper snow cover at sites increasingly exposed to frost due to forest loss and weakened microclimatic buffering (Örlander, 1993; Simons et al., 2010). Such a connection of baseline snow cover with the impacts of forest loss could indicate potential future effects of climate change: greater decreases in snow depth may have an exacerbating effect on the extinction probability of *C. bulbosa* together with land use change. Thus, although our data did not reveal an effect of temporal change in snow cover on extinction probability, we expect that accumulating data with proceeding climate change may come to identify such effects. Our results, together with e.g. Aalto et al. (2023) show that on average, snow depth is decreasing in many areas in Finland, although there is regional variation in the direction of the trend. Thus, declining snow depth appears to be an ongoing component of climate change in Finland, likely posing increasing pressures on species that rely on snow.

### 4.3 Separate effects of climate were weak compared to the stronger interactive effects of climate with forest loss

In addition to the interactive effects of land use change and climate, we found some indications of separate effects of climate. Baseline GDD5 and temporal change in GDD5 were both associated with extinction probability, although the effects were relatively statistically weak. Extinction probability was somewhat higher at sites with higher baseline GDD5 and in sites with greater decrease in GDD5. Moreover, it should be acknowledged that baseline thermal accumulation showed moderate multicollinearity, due to which its effects should be inferred with caution. Notwithstanding, such opposing associations of baseline GDD5 and its temporal change could be explained by increased drought effects at sites with higher baseline thermal accumulation (Wu et al., 2025; Yan et al., 2025), whereas greater decrease in thermal accumulation may negatively influence survival, e.g., through a shorter growing season.

### 4.4 The current impact of forest loss originated from local scale but landscape scale forest loss may also matter

We only found significant effects of forest loss on the extinction probability of *C. bulbosa* at the local, rather than the landscape scale. This result is in line with the general understanding that the effects of habitat loss, fragmentation and degradation are scale-dependent (Banks-Leite et al., 2020). This scale-dependent effect of forest loss suggests that currently, forest loss is primarily influencing the persistence of *C. bulbosa* through the loss of suitable microsites (Eriksson & Ehrlén, 1992). However, in the long run, forest loss could also influence the species at the landscape-scale through seed limitation (Eriksson & Ehrlén, 1992). This is because landscape-scale forest loss may lead to the loss of neighboring populations, and thus connectivity of the remaining populations. Reduced connectivity, in turn, can influence the persistence of populations through reducing seed exchange between populations and recolonization of suitable habitats (Moilanen & Hanski, 2001). This kind of a negative effect of connectivity reduction on the persistence of *C. bulbosa* was supported by our intermediate models (i.e. those that did not include interactions), showing that higher connectivity decreased the extinction probability of *C. bulbosa*. In addition to this effect, we also observed an interactive effect between baseline snow conditions and connectivity (such that extinction probability was greatest in well-connected sites with deep baseline snow cover). However, this effect is likely a spurious finding driven by a lack of observations at the extreme ends of baseline snow depth. While interactions of connectivity with land use and climate change remain inconclusive, our data indicates that connectivity can independently increase the probability of the species to persist.

### 4.5 Additional factors that may explain extinction probability

Overall, our results highlight the importance of land use change and climate in explaining variation in the extinction probability of *C. bulbosa*. At the same time, the results indicate that several factors not considered in our analyses are likely to play a role in explaining the extinction probability of this species. One such potentially important factor could be soil composition, as *C. bulbosa* is known to favor soils with high calcium concentrations (Hyvärinen et al. 2019). Others include factors connected to biotic interactions with pollinators and symbiotic partners, as suggested by a previous analysis of the drivers of occurrence and abundance patterns of *C. bulbosa* (Koivusaari et al., 2026). Finally, while we focused on climatic conditions during and preceding the flowering season of *C. bulbosa*, conditions during other seasons may also impact the species’ performance and viability. For example, autumn leaf production and winter dormancy may affect its persistence over the dormant period and thus its regenerative potential in spring. Future studies should assess the effects of conditions throughout the year, interactions with other species, soil conditions and their changes over time when assessing the extinction probability of *C. bulbosa*.

## 5. CONCLUSIONS

Land use and climate change are two of the greatest threats to biodiversity globally, but the mechanisms by which they affect species persistence in boreal forests, a globally vital ecosystem, remain poorly understood. Our results show that land use change, baseline climate and climate change interact strongly in shaping extinction risk for a boreal forest orchid, jointly explaining 57% of the variation in extinction probability. We also find that the effects of forest loss are scale-dependent, originating mainly from the local rather than the landscape-scale. These findings highlight that land use and climate change do not act in isolation in driving biodiversity responses, and emphasizes the need to consider their interactions when predicting species’ persistence under environmental change.

## CONFLICT OF INTEREST STATEMENT

The authors declare no conflicts of interest.

## ACKNOWLEDGEMENTS

SHMK, ALKM, TL and ML acknowledge funding from the Kone Foundation, SHMK, RKH, and MHH from the Research Council of Finland (grant no. 360742 to MHH), and SHMK additionally from Societas pro Fauna et Flora Fennica, Waldemar von Frenckells Stiftelse, and Nordenskiöld-Samfundet.

## AUTHOR CONTRIBUTIONS

**S. H. M. Koivusaari:** conceptualization (lead), data curation (lead), formal analysis (lead), funding acquisition (equal), investigation (lead), methodology (lead), project administration (lead), resources (lead), software (lead), validation (lead), writing – original draft (lead), writing – review & editing (lead). **M. Luoto:** conceptualization (equal), formal analysis (equal), funding acquisition (equal), methodology (equal), project administration (equal), resources (supporting), supervision (lead), validation (equal), writing – original draft (equal), writing – review & editing (equal). **J. Aalto:** conceptualization (supporting), writing – review and editing (supporting). **K. Aapala:** conceptualization (equal), writing – review and editing (supporting). **M. H. Hällfors:** conceptualization (supporting), formal analysis (supporting), funding acquisition (equal), methodology (equal), project administration (equal), resources (supporting), supervision (equal), validation (supporting), writing – original draft (equal), writing – review & editing (equal). **R. K. Heikkinen:** conceptualization (equal), writing – review and editing (supporting). **T. Laurinen:** investigation (supporting), writing – review and editing (supporting). **A. L. K. Mattila:** conceptualization (equal), data curation (equal), formal analysis (supporting), funding acquisition (equal), investigation (equal), methodology (equal), project administration (equal), resources (supporting), supervision (equal), validation (supporting), writing – original draft (equal), writing – review & editing (equal).

## Notes

### Competing Interest Statement

The authors have declared no competing interest.

## REFERENCES

Aalto, J., Kämäräinen, M., Rantanen, M., Niittynen, P., Phoenix, G. K., Lenoir, J., Maclean, I., & Luoto, M. (2026). A new era of bioclimatic extremes in the terrestrial Arctic. Science Advances, 12(2), eadw5698. 10.1126/sciadv.adw5698

Aalto, J., Lehtonen, I., Pirinen, P., Aapala, K., & Heikkinen, R. K. (2023). Bioclimate change across the protected area network of Finland. Science of The Total Environment, 893, 164782. 10.1016/j.scitotenv.2023.164782

Aalto, J., Pirinen, P., & Jylhä, K. (2016). New gridded daily climatology of Finland: Permutation-based uncertainty estimates and temporal trends in climate. Journal of Geophysical Research: Atmospheres, 121(8), 3807–3823. 10.1002/2015JD024651

Abeli, T., Jäkäläniemi, A., Wannas, L., Mutikainen, P., & Tuomi, J. (2013). Pollen limitation and fruiting failure related to canopy closure in Calypso bulbosa (Orchidaceae), a northern food-deceptive orchid with a single flower. Botanical Journal of the Linnean Society, 171(4), 744–750. 10.1111/boj.12014

Ackerman, J. D. (1981). Pollination Biology of Calypso Bulbosa Var. Occidentals (orchidaceae): A Food-Deception System. Madroño, 28(3), 101–110.

Alexandersson, R., & Ågren, J. (1996). Population size, pollinator visitation and fruit production in the deceptive orchid Calypso bulbosa. Oecologia, 107(4), 533–540. 10.1007/BF00333945

Alexandersson, R., & Ågren, J. (2000). Genetic structure in the nonrewarding, bumblebee-pollinated orchid Calypso bulbosa. Heredity, 85(4), 401–409. 10.1046/j.1365-2540.2000.00777.x

Auffret, A. G., & Svenning, J.-C. (2022). Climate warming has compounded plant responses to habitat conversion in northern Europe. Nature Communications, 13(1), 7818. 10.1038/s41467-022-35516-7

Banks-Leite, C., Ewers, R. M., Folkard-Tapp, H., & Fraser, A. (2020). Countering the effects of habitat loss, fragmentation, and degradation through habitat restoration. One Earth, 3(6), 672–676. 10.1016/j.oneear.2020.11.016

Boyden, T. C. (1982). The pollination biology of Calypso bulbosa var. Americana (Orchidaceae): Initial deception of bumblebee visitors. Oecologia, 55(2), 178–184. 10.1007/BF00384485

Brown, J. H., & Kodric-Brown, A. (1977). Turnover Rates in Insular Biogeography: Effect of Immigration on Extinction. Ecology, 58(2), 445–449. 10.2307/1935620

Burton P. J., Bergeron Y., Bogdanski, B. E. C., et al. (2010). Sustainability of boreal forests and forestry in a changing environment. In G. Mery, P. Katila, G. Galloway, R. I. Alfaro, M. Kanninen, M. Lobovikov, J. Varjo (Eds.), Forests and society—responding to global drivers of change (pp. 249–282). International Union of Forest Research Organizations, IUFRO.

Chen, J., Franklin, J. F., & Spies, T. A. (1993). Contrasting microclimates among clearcut, edge, and interior of old-growth Douglas-fir forest. Agricultural and Forest Meteorology, 63(3), 219–237. 10.1016/0168-1923(93)90061-L

Currah, R. S., Hambleton, S., & Smreciu, A. (1988). MYCORRHIZAE AND MYCORRHIZAL FUNGI OF CALYPSO BULBOSA. American Journal of Botany, 75(5), 739–752. 10.1002/j.1537-2197.1988.tb13495.x

De Frenne, P., Zellweger, F., Rodríguez-Sánchez, F., Scheffers, B. R., Hylander, K., Luoto, M., Vellend, M., Verheyen, K., & Lenoir, J. (2019). Global buffering of temperatures under forest canopies. Nature Ecology & Evolution, 3(5), 744–749. 10.1038/s41559-019-0842-1

Deutsch, C. A., Tewksbury, J. J., Huey, R. B., Sheldon, K. S., Ghalambor, C. K., Haak, D. C., & Martin, P. R. (2008). Impacts of climate warming on terrestrial ectotherms across latitude. Proceedings of the National Academy of Sciences of the United States of America, 105(18), 6668–6672. 10.1073/pnas.0709472105

Eriksson, O., & Ehrlén, J. (1992). Seed and microsite limitation of recruitment in plant populations. Oecologia, 91(3), 360–364. 10.1007/BF00317624

Finnish Biodiversity Information Facility (FinBIF) (2023). Browse & Search. Retrieved February 23, 2023, from https://laji.fi/en/observation/list.

Frey, S. J. K., Hadley, A. S., Johnson, S. L., Schulze, M., Jones, J. A., & Betts, M. G. (2016). Spatial models reveal the microclimatic buffering capacity of old-growth forests. Science Advances, 2(4), e1501392. 10.1126/sciadv.1501392

Gauthier, S., Bernier, P., Kuuluvainen, T., Shvidenko, A. Z., & Schepaschenko, D. G. (2015). Boreal forest health and global change. Science, 349(6250), 819–822. 10.1126/science.aaa9092

Gauthier, S., Kuuluvainen, T., Macdonald, S. E., Shorohova, E., Shvidenko, A., Bélisle, A.-C., Vaillancourt, M.-A., Leduc, A., Grosbois, G., Bergeron, Y., Morin, H., Girona, M. M. (2023). Ecosystem Management of the Boreal Forest in the Era of Global Change. Girona, M. M., Morin, H., Gauthier, S. (eds.) (2023). Boreal Forests in the Face of Climate Change. Advances in Global Change Research 74.

Geiger, R. (1965). The Climate Near the Ground. Harvard University Press.

Hadley, G. (1982). Orchid mycorrhiza. In J. Arditti [ed.], Orchid biology: reviews and perspectives II, 82-118. Comell University Press, Ithaca, NY.

Hanski, I. (1994). A Practical Model of Metapopulation Dynamics. The Journal of Animal Ecology, 63(1), 151. 10.2307/5591

Hartig, F. (2024). _DHARMa: Residual Diagnostics for Hierarchical (Multi-Level / Mixed) Regression Models_. doi:10.32614/CRAN.package.DHARMa <10.32614/CRAN.package.DHARMa>, R package version 0.4.7, <https://CRAN.R-project.org/package=DHARMa>.

Hijmans R (2025). _terra: Spatial Data Analysis_. doi:10.32614/CRAN.package.terra <10.32614/CRAN.package.terra>, R package version 1.8-70, <https://CRAN.R-project.org/package=terra>.

Hirabayashi, Y., Mahendran, R., Koirala, S., Konoshima, L., Yamazaki, D., Watanabe, S., Kim, H., & Kanae, S. (2013). Global flood risk under climate change. Nature Climate Change, 3(9), 816–821. 10.1038/nclimate1911

Hu, J., Zhao, X., Gu, L., Liu, P., Zhao, B., Zhang, J., & Ren, B. (2023). The effects of high temperature, drought, and their combined stresses on the photosynthesis and senescence of summer maize. Agricultural Water Management, 289, 108525. 10.1016/j.agwat.2023.108525

Hyvärinen, E., Juslén, A., Kemppainen, E., Uddström, A. & Liukko, U.-M. (eds.) (2019). The 2019 Red List of Finnish Species. Ympäristöministeriö & Suomen ympäristökeskus. Helsinki. 704 p.

IPBES (2019). Summary for policymakers of the global assessment report on biodiversity and ecosystem services of the Intergovernmental Science-Policy Platform on Biodiversity and Ecosystem Services. S. Díaz, J. Settele, E. S. Brondízio >E.S., H. T. Ngo, M. Guèze, J. Agard, A. Arneth, P. Balvanera, K. A. Brauman, S. H. M. Butchart, K. M. A. Chan, L. A. Garibaldi, K. Ichii, J. Liu, S. M. Subramanian, G. F. Midgley, P. Miloslavich, Z. Molnár, D. Obura, A. Pfaff, S. Polasky, A. Purvis, J. Razzaque, B. Reyers, R. Roy Chowdhury, Y. J. Shin, I. J. Visseren-Hamakers, K. J. Willis, and C. N. Zayas (eds.). IPBES secretariat, Bonn, Germany. 56 pages

Irfan, M., Hayat, S., Hayat, Q., Afroz, S., & Ahmad, A. (2010). Physiological and biochemical changes in plants under waterlogging. Protoplasma, 241(1), 3–17. 10.1007/s00709-009-0098-8

Jaureguiberry, P., Titeux, N., Wiemers, M., Bowler, D. E., Coscieme, L., Golden, A. S., Guerra, C. A., Jacob, U., Takahashi, Y., Settele, J., Díaz, S., Molnár, Z., & Purvis, A. (2022). The direct drivers of recent global anthropogenic biodiversity loss. Science Advances, 8(45), eabm9982. 10.1126/sciadv.abm9982

Kemppinen, J., Lembrechts, J. J., Van Meerbeek, K., Carnicer, J., Chardon, N. I., Kardol, P., Lenoir, J., Liu, D., Maclean, I., Pergl, J., Saccone, P., Senior, R. A., Shen, T., Słowińska, S., Vandvik, V., von Oppen, J., Aalto, J., Ayalew, B., Bates, O., … De Frenne, P. (2024). Microclimate, an important part of ecology and biogeography. Global Ecology and Biogeography, 33(6), e13834. 10.1111/geb.13834

Koivusaari, S., Mattila, A., Hällfors, M., Aalto, J., Aapala, K., Heikkinen, R., Kemppinen, J., Laurinen, T., Lehtinen, J., Mostarin, M., Niittynen, P., Opedal, Ø., & Luoto, M. (n.d.). Forest features shape the occurrence and abundance of a threatened boreal orchid Calypso bulbosa via microclimatic conditions and vegetation composition. Authorea, 2026(0727). 10.22541/authorea.15006583/v1

Lososová, Z., Axmanová, I., Chytrý, M., Midolo, G., Abdulhak, S., Karger, D. N., Renaud, J., Van Es, J., Vittoz, P., & Thuiller, W. (2023). Seed dispersal distance classes and dispersal modes for the European flora. Global Ecology and Biogeography, 32(9), 1485–1494. 10.1111/geb.13712

Lüdecke, D. (2018). “ggeffects: Tidy Data Frames of Marginal Effects from Regression Models.” _Journal of Open Source Software_, *3*(26), 772. doi:10.21105/joss.00772 <10.21105/joss.00772>.

Martin, A. E., Lockhart, J. K., & Fahrig, L. (n.d.). Are weak dispersers more vulnerable than strong dispersers to land use intensification? Proceedings of the Royal Society B: Biological Sciences, 290(1990), 20220909. 10.1098/rspb.2022.0909

Mattila, A. L. K., Opedal, Ø. H., Hällfors, M. H., Pietikäinen, L., Koivusaari, S. H. M., & Hyvärinen, M.-T. (2024). The potential for evolutionary rescue in an Arctic seashore plant threatened by climate change. Proceedings of the Royal Society B: Biological Sciences, 291(2032), 20241351. 10.1098/rspb.2024.1351

Meeussen, C., Govaert, S., Vanneste, T., Haesen, S., Van Meerbeek, K., Bollmann, K., Brunet, J., Calders, K., Cousins, S. A. O., Diekmann, M., Graae, B. J., Iacopetti, G., Lenoir, J.,Orczewska, A., Ponette, Q., Plue, J., Selvi, F., Spicher, F., Sorensen, M. V., … De Frenne, P. (2021). Drivers of carbon stocks in forest edges across Europe. Science of the Total Environment, 759(143497). 10.1016/j.scitotenv.2020.143497

Menge, J. H., Magdon, P., Wöllauer, S., & Ehbrecht, M. (2023). Impacts of forest management on stand and landscape-level microclimate heterogeneity of European beech forests. Landscape Ecology, 38(4), 903–917. 10.1007/s10980-023-01596-z

Moilanen, A., & Hanski, I. (2001). On the use of connectivity measures in spatial ecology. Oikos, 95(1), 147–151.

Mosquin, T. (1970). The Reproductive Biology of Calypso bulbosa (Orchidaceae). The Canadian Field-Naturalist, 84(3), 291–296. 10.5962/p.342973

Northrup, J. M., Rivers, J. W., Yang, Z., & Betts, M. G. (2019). Synergistic effects of climate and land-use change influence broad-scale avian population declines. Global Change Biology, 25(5), 1561–1575. 10.1111/gcb.14571

Oliver, T. H., & Morecroft, M. D. (2014). Interactions between climate change and land use change on biodiversity: Attribution problems, risks, and opportunities. Wiley Interdisciplinary Reviews: Climate Change, 5(3), 317–335.

Örlander, G. (1993). Shading Reduces both Visible and Invisible Frost Damage to Norway Spruce Seedlings in the Field. Forestry: An International Journal of Forest Research, 66(1), 27–36. 10.1093/forestry/66.1.27

Pohlert, T. (2023). _trend: Non-Parametric Trend Tests and Change-Point Detection_. doi:10.32614/CRAN.package.trend <10.32614/CRAN.package.trend>, R package version 1.1.6, <https://CRAN.R-project.org/package=trend>.

Rao, M.P., Davi, N.K., Magney, T.S. et al. Approaching a thermal tipping point in the Eurasian boreal forest at its southern margin. Commun Earth Environ 4, 247 (2023). 10.1038/s43247-023-00910-6

Rantanen, M., Karpechko, A. Y., Lipponen, A., Nordling, K., Hyvärinen, O., Ruosteenoja, K., Vihma, T., & Laaksonen, A. (2022). The Arctic has warmed nearly four times faster than the globe since 1979. Communications Earth & Environment, 3(1), 168. 10.1038/s43247-022-00498-3

R Core Team (2025). _R: A Language and Environment for Statistical Computing_. R Foundation for Statistical Computing, Vienna, Austria. <https://www.R-project.org/>.

Robin, X., Turck, N., Hainard, A., Tiberti, N., Lisacek, F., Sanchez, J. C. & Müller, M. (2011). pROC: an open-source package for R and S+ to analyze and compare ROC curves. BMC Bioinformatics, 12, p. 77. DOI: 10.1186/1471-2105-12-77 <http://www.biomedcentral.com/1471-2105/12/77/>

Román-Palacios, C., & Wiens, J. J. (2020). Recent responses to climate change reveal the drivers of species extinction and survival. Proceedings of the National Academy of Sciences, 117(8), 4211–4217. 10.1073/pnas.1913007117

Saccheri, I., Kuussaari, M., Kankare, M., Vikman, P., Fortelius, W., & Hanski, I. (1998). Inbreeding and extinction in a butterfly metapopulation. Nature, 392(6675), 491–494. 10.1038/33136

Semenchuk, P., Plutzar, C., Kastner, T., Matej, S., Bidoglio, G., Erb, K.-H., Essl, F., Haberl, H., Wessely, J., Krausmann, F., & Dullinger, S. (2022). Relative effects of land conversion and land-use intensity on terrestrial vertebrate diversity. Nature Communications, 13(1), 615. 10.1038/s41467-022-28245-4

Sen, P. K. (1968). Estimates of the Regression Coefficient Based on Kendall’s Tau. Journal of the American Statistical Association, 63(324), 1379–1389. 10.2307/2285891

Senior, R. A., Hill, J. K., González del Pliego, P., Goode, L. K., & Edwards, D. P. (2017). A pantropical analysis of the impacts of forest degradation and conversion on local temperature. Ecology and Evolution, 7(19), 7897–7908. 10.1002/ece3.3262

Simons, A. M., Goulet, J. M., & Bellehumeur, K. F. (2010). The effect of snow depth on overwinter survival in Lobelia inflata. Oikos, 119(10), 1685–1689. 10.1111/j.1600-0706.2010.18515.x

Starck, I., Aalto, J., Hancock, S., Kemppinen, J., Lehtinen, J., Luoto, M., Niittynen, P., Rissanen, T., Tyystjärvi, V., Valkonen, S., & Maeda, E. (2026). Limited microclimatic buffering capacity in boreal forests calls for sustainable management strategies. Environmental Research Letters, 21(1), 014011. 10.1088/1748-9326/ae2d79

Starck, I., Aalto, J., Hancock, S., Valkonen, S., Kalliovirta, L., & Maeda, E. (2025). Slow recovery of microclimate temperature buffering capacity after clear-cuts in boreal forests. Agricultural and Forest Meteorology, 363, 110434. 10.1016/j.agrformet.2025.110434

Suppula, M., Hällfors, M. H., Aapala, K., Aalto, J., Kemppainen, E., Leikola, N., Pirinen, P., & Heikkinen, R. K. (2023). Climate and land-use change drive population decline in a red-listed plant species. Global Ecology and Conservation, 45, e02526. 10.1016/j.gecco.2023.e02526

Thomas, C. D., Cameron, A., Green, R. E., Bakkenes, M., Beaumont, L. J., Collingham, Y. C., Erasmus, B. F. N., de Siqueira, M. F., Grainger, A., Hannah, L., Hughes, L., Huntley, B., van Jaarsveld, A. S., Midgley, G. F., Miles, L., Ortega-Huerta, M. A., Townsend Peterson, A., Phillips, O. L., & Williams, S. E. (2004). Extinction risk from climate change. Nature, 427(6970), 145–148. 10.1038/nature02121

Tuomi, J., Lämsä, J., Wannas, L., Abeli, T., & Jäkäläniemi, A. (2015). Pollinator Behaviour on a Food-Deceptive Orchid Calypso bulbosa and Coflowering Species. The Scientific World Journal, 2015(1), 482161. 10.1155/2015/482161

Wu, H., Su, X., Huang, S., Singh, V. P., Zhou, S., Tan, X., & Hu, X. (2025). Decreasing dynamic predictability of global agricultural drought with warming climate. Nature Climate Change, 15(4), 411–419. 10.1038/s41558-025-02289-y

Yan, H., Sun, N., Yao, L., Thurber, T. B., & Rice, J. S. (2025). Rising temperatures intensify drought propagation and severity across the contiguous United States. Npj Natural Hazards, 2(1), 91. 10.1038/s44304-025-00134-y

Zellweger, F., De Frenne, P., Lenoir, J., Vangansbeke, P., Verheyen, K., Bernhardt-Römermann, M., Baeten, L., Hédl, R., Berki, I., Brunet, J., Van Calster, H., Chudomelová, M., Decocq, G., Dirnböck, T., Durak, T., Heinken, T., Jaroszewicz, B., Kopecký, M., Máliš, F., … Coomes, D. (2020). Forest microclimate dynamics drive plant responses to warming. Science, 368(6492), 772–775. 10.1126/science.aba6880

Zhang, Y., Chen, X., Geng, S., & Zhang, X. (2025). A review of soil waterlogging impacts, mechanisms, and adaptive strategies. Frontiers in Plant Science, 16, 1545912. 10.3389/fpls.2025.1545912

