## Supplementary information for "Interactive effects of land use change and climate drive the decline of a threatened boreal forest orchid in Finland"

**Contents:**

Figures S1-S2
Table S1

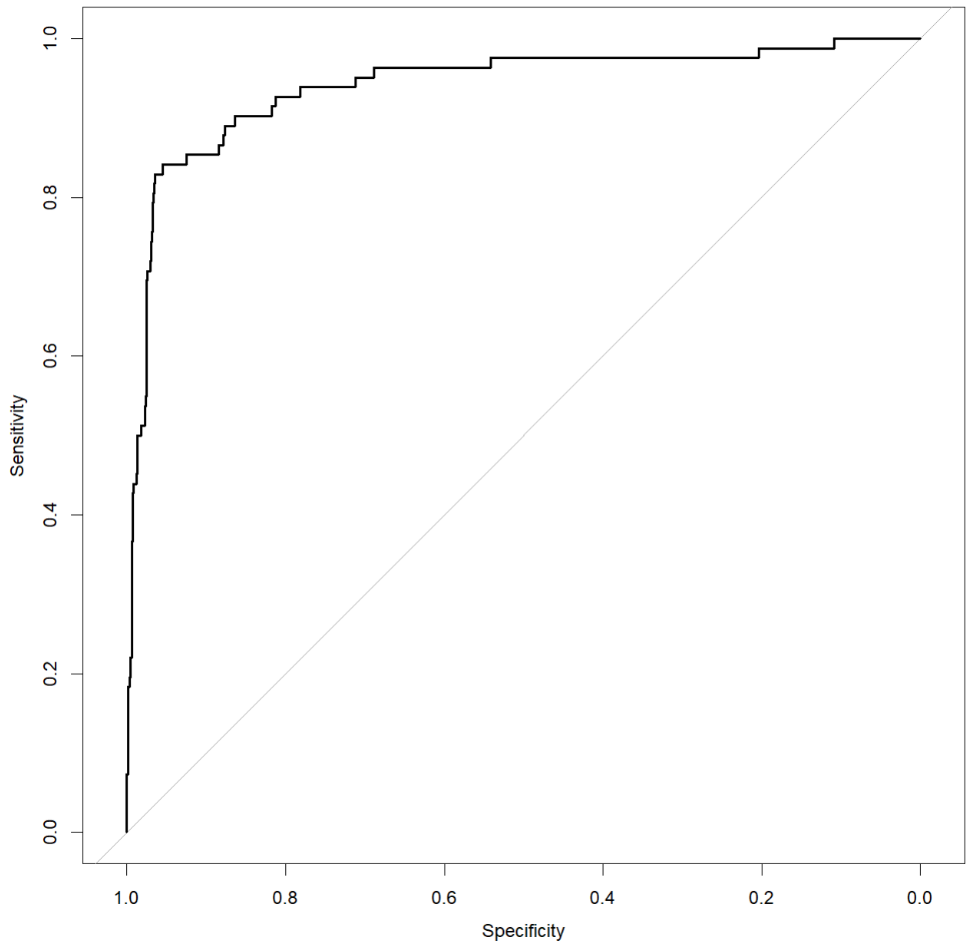


**Figure S1.** ROC-curve for the best performing (lowest AIC and highest R2) model i.e. the model 4 that included land use change, climate, connectivity, and significant interactions between these. AUC (Area under the curve) = 0.94.

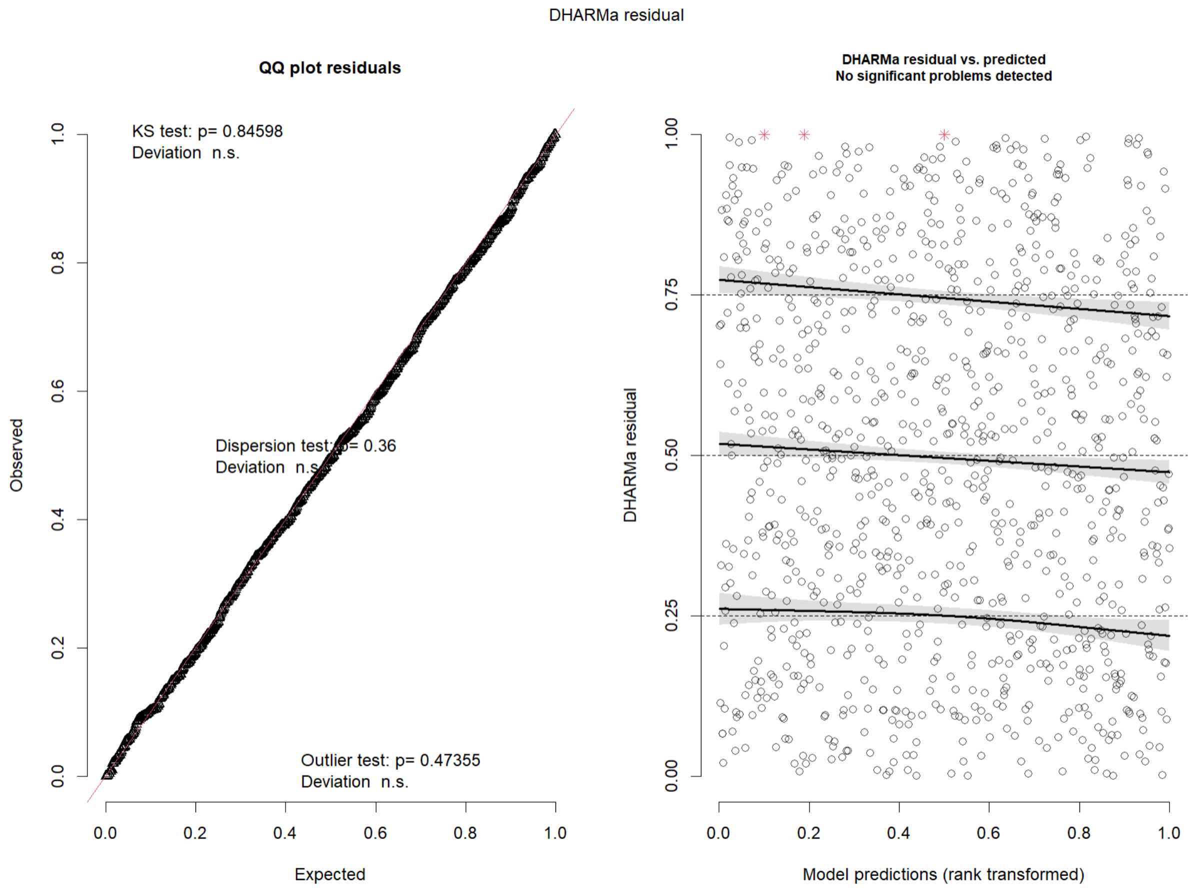


**Figure S2.** QQ-plots and residuals vs. fitted values plots for the best performing (lowest AIC and highest R2) model i.e. the model 4 that included land use change, climate, connectivity, and significant interactions between these.

**Table S1.** Summary statistics for models fitted using connectivity data calculated with method 1 (random mean abundance values for localities with NA or 0 mean abundance). CMP = change in mature forest presence, CMC = change in mature forest cover, CRF = change in rainfall, CGDD = change in GDD5, CSD = change in snow depth, MRF = mean rainfall, MGDD = mean GDD5, MSD = mean snow depth, CON = connectivity.

| Model | Term | Estimate | Std. Error | Statistic | p-value |
|---|---|---|---|---|---|
| LUC | (Intercept) | 0.71 | 0.41 | 1.74 | 0.083 |
| | CMP | -1.23 | 0.13 | -9.39 | 0.000 |
| | CMC | 0.00 | 0.00 | 0.19 | 0.851 |
| | CON | 0.01 | 0.01 | 1.25 | 0.212 |
| CC | (Intercept) | 7.27 | 3.09 | 2.36 | 0.018 |
| | CRF | -12.05 | 1.68 | -7.17 | 0.000 |
| | CGDD | -0.53 | 0.46 | -1.15 | 0.249 |
| | CSD | 3.04 | 1.87 | 1.62 | 0.105 |
| | MRF | -0.26 | 0.06 | -4.64 | 0.000 |
| | MGDD | 0.01 | 0.01 | 1.09 | 0.275 |
| | MSD | 0.02 | 0.04 | 0.54 | 0.591 |
| | CON | -0.01 | 0.01 | -2.42 | 0.015 |
| LUC + CC | (Intercept) | 7.02 | 4.05 | 1.73 | 0.083 |
| | CMP | -1.43 | 0.17 | -8.52 | 0.000 |
| | CMC | 0.00 | 0.00 | -0.05 | 0.963 |
| | CRF | -15.21 | 2.15 | -7.07 | 0.000 |
| | CGDD | 0.04 | 0.50 | 0.07 | 0.944 |
| | CSD | 3.72 | 1.86 | 2.00 | 0.045 |
| | MRF | -0.10 | 0.07 | -1.4 | 0.161 |
| | MGDD | 0.01 | 0.01 | 0.74 | 0.461 |
| | MSD | -0.03 | 0.06 | -0.58 | 0.56 |
| | CON | 0.00 | 0.01 | 0.35 | 0.728 |
| LUC + CC + LUC:CC | (Intercept) | -37.77 | 11.14 | -3.39 | 0.001 |
| | CMP | 8.64 | 2.59 | 3.34 | 0.001 |
| | CMC | 0.00 | 0.01 | -0.01 | 0.994 |
| | CRF | 0.11 | 4.57 | 0.02 | 0.980 |
| | CGDD | -0.75 | 0.57 | -1.30 | 0.193 |
| | CSD | 7.75 | 2.72 | 2.85 | 0.004 |
| | MRF | 0.88 | 0.27 | 3.22 | 0.001 |
| | MGDD | 0.05 | 0.02 | 2.82 | 0.005 |
| | MSD | -0.22 | 0.12 | -1.79 | 0.074 |
| | CON | 0.98 | 0.33 | 3.01 | 0.003 |
| | CMP:CRF | -8.54 | 2.12 | -4.04 | 0.000 |
| | CMP:MRF | -0.29 | 0.09 | -3.42 | 0.001 |
| | CMP:MSD | 0.08 | 0.04 | 1.98 | 0.047 |
| | CMP:CON | -0.02 | 0.01 | -2.86 | 0.004 |
| | CSD:CON | -0.28 | 0.12 | -2.39 | 0.017 |
| | MRF:CON | -0.01 | 0.01 | -2.68 | 0.007 |
| | MGDD:CON | 0.00 | 0.00 | -2.47 | 0.014 |
